# Efficient and scalable prediction of spatio-temporal stochastic gene expression in cells and tissues using graph neural networks

**DOI:** 10.1101/2023.02.28.530379

**Authors:** Zhixing Cao, Rui Chen, Libin Xu, Xinyi Zhou, Xiaoming Fu, Weimin Zhong, Ramon Grima

## Abstract

The simulation of spatial stochastic models is highly computationally expensive, an issue that has severely limited our understanding of the spatial nature of gene expression. Here we devise a graph neural network based method to learn, from stochastic trajectories in a small region of space, an effective master equation for the time-dependent marginal probability distributions of mRNA and protein numbers at sub-cellular resolution for every cell in a tissue. Numerical solution of this equation leads to accurate results in a small fraction of the computation time of standard simulation methods. Moreover its predictions can be extrapolated to a spatial organisation (a cell network topology) and regions of parameter space unseen in its neural network training. The scalability and accuracy of the method suggest it is a promising approach for whole cell modelling and for detailed comparisons of stochastic models with spatial genomics data.

## Introduction

In the past two decades, experiments have firmly established the stochastic nature of gene expression [1–4]. Our understanding of gene product dynamics has been largely helped by the development of mathematical models of gene expression [5–10]. Analytical and simulation-based studies have elucidated how cells control and exploit noise for optimal biological functioning [11,12] and have been used to fit data to estimate kinetic parameter values [13–17]. However the vast literature on this subject has been largely focused on understanding molecular fluctuations at the single cell level, e.g. the fluctuations of the total number of mRNA in a cell. But this is certainly not the only spatial scale at which molecular fluctuations are important – it is of increasing interest to understand these fluctuations at finer spatial scales, e.g. spatio-temporal patterns of mRNA and protein across intracellular space reflecting their transport and localisation properties. Moreover it is of significant interest to understand molecular fluctuations in all cells in a tissue simultaneously since their spatial organisation influences biological function [18–22].

While single-cell models avoid any description of space by implicitly assuming well-mixing, the construction of simulations at the intra-cellular and super-cellular scales necessitates this additional ingredient. Herein lies a major stumbling block because the computational cost of spatial stochastic simulation using conventional techniques can be enormous. In this case, typically space is discretised into a finite number of volumes (voxels) and rules are defined to model reactions occurring within each voxel and molecular transport across neighbouring voxels. Applying a conventional simulation technique, Finite State Projection (FSP [23]), to the reaction-diffusion master equation (RDME [24–27]) corresponding to this set of processes implies solving *N^MS^* coupled differential equations (one for each state of the system) where *N* – 1 is a user-defined upper bound on the molecule counts for all species in each voxel, *M* is the number of voxels and *S* is the number of species. The use of Monte Carlo techniques such as Gillespie’s stochastic simulation algorithm (SSA [28–32]) avoids the solution of these differential equations, but the computation time can still be very significant because (i) the effective number of species and the number of reactions increase linearly with *M* and (ii) a considerably large amount of ensemble averaging of the trajectories may be needed to obtain statistically accurate results for the distributions of molecular counts in each voxel. This makes the practical exploration of a spatially extended system’s properties across parameter space very challenging.

In this article, we devise an alternative method which uses graph neural networks (GNNs) to circumvent the aforementioned issues. A GNN is a type of neural network structure operating on graph-structured data which has found several applications in RNA-seq data analyses [33,34], biomedicine [35] and material science [36]. It has powerful capabilities of extrapolation to different datasets and topologies, and avoids overfitting [37,38]. Here we utilise GNNs to learn, from a small sample of SSA simulations, the propensity functions of an effective master equation for the marginal distributions of each species in each voxel. Application of FSP to the latter implies the solution of *MNS* equations, typically a very small fraction of the *N^MS^* equations given by the conventional FSP. In addition, the method’s ability to extrapolate predictions to voxel network topologies and regions of parameter space unseen in its training gives it a distinctive computational advantage over the SSA.

## Results

### Graph neural networks provide an accurate and computationally cheap approximation of the RDME: illustration on a single-species system

We illustrate our novel approach by considering a system of *m* cells with biochemical reactions occurring inside them and the transport of molecules between them. Specifically in each cell *i*, we model the production of proteins by a zero-order reaction with rate *ρ_i_* and their degradation/dilution by a first-order reaction with rate *d_i_*. Additionally, we model the active transport of protein molecules from cell *i* to a neighbouring cell *j* by a first-order reaction with a Michaelis-Menten propensity 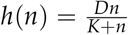 where the saturation at large *n* stems from the finite maximum speed of the transporter molecules. This is an elementary model of a tissue [19]. The reaction scheme describing this system is given by:

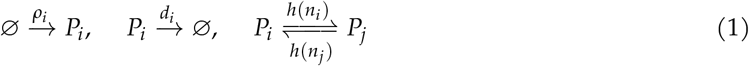

for *i* = 1,…, *m* and 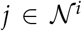 where 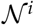 is the set of neighbours of cell *i* which effectively encodes the topology of cell-cell interactions. The multi-cell system and the associated reactions are also illustrated in Fig. 1a. Assuming Markovian dynamics, the stochastic dynamics are described by the reaction-diffusion master equation:

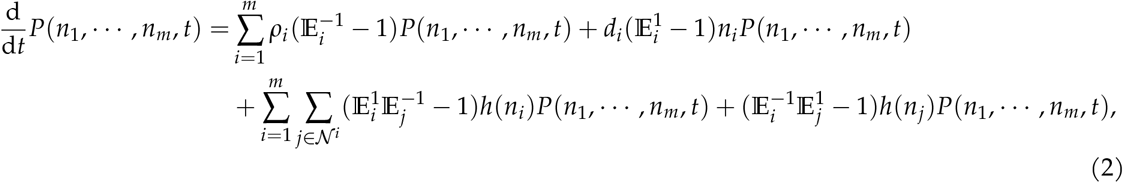

where *P*(*n*_1_,…, *n_m_, t*) is the probability of observing at time *t*, cell 1 with *n*_1_ protein molecules, cell 2 with *n*_2_ protein molecules, …, cell *m* with *n_m_* protein molecules. The operator 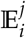 is a step operator defined as 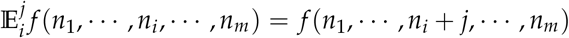. If we wanted to solve Eq. (2) using the FSP algorithm, on a state space truncated up to a maximum molecule number *N* – 1, then we would need to integrate *N^m^* coupled differential equations. It is clear that for very modest maximum protein numbers, say a few tens of molecules, this algorithm is computationally prohibitive if we are interested in studying a system with more than a handful of interconnected cells.

**Figure 1:**
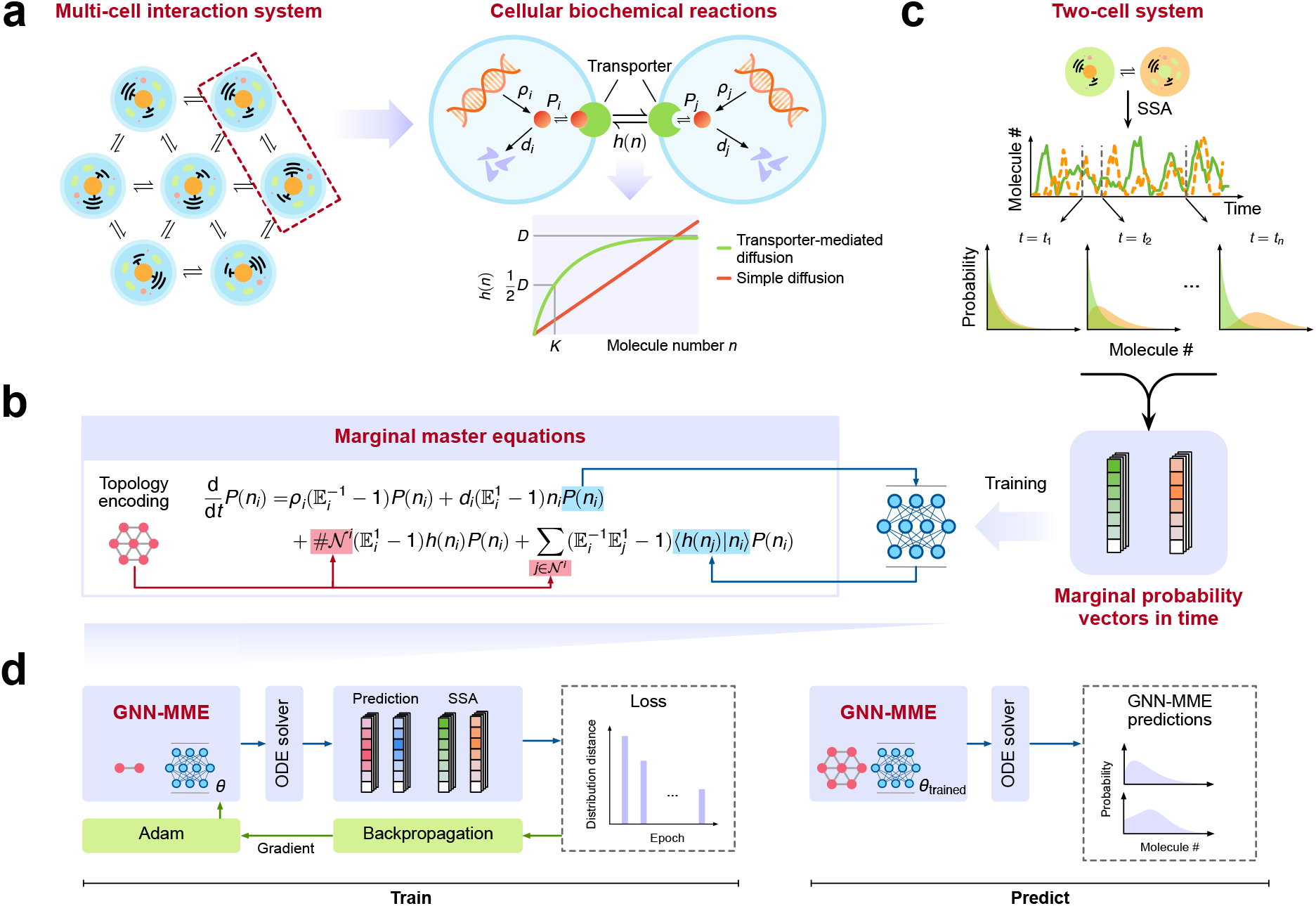
Graph neural network aided construction of master equations for the marginal distributions of protein counts in a multi-cell system. (a) Illustration of a connected system of *m* cells – proteins (solid red circles) are produced and degraded in cell *i* with rates *p_i_* and *d_i_*, respectively and they are transported between cells by membrane transport proteins (transporters) such as carrier proteins (green pac-mans). The propensity function modelling active transport *h*(*n*) is a Michaelis-Menten function of the protein number which effectively captures the saturation of carrier proteins at high number. (b) From the master equation of the multi-cell system, a master equation for the marginal distributions is derived (Eq. (3)). The terms describing transport influx into a cell are unknown functions of the marginal distributions. Hence the master equations are not closed and cannot be solved. We use a neural network, trained using SSA simulations of two-cell systems (c), to learn the function that maps the marginal distributions to the effective propensity describing influx into a cell, thus closing the master equations – we call the latter a *graph neural network modularised master equation* (*GNN-MME*). (d) Left panel. An optimisation procedure is used to find the neural network coefficients which minimise the distance between the FSP solution of the two-cell GNN-MME and the SSA marginal distributions. This constitutes the learning of the functional approximation of the effective influx propensity by the neural network. (d) Right panel. The learnt propensities from two-cell simulations are used to directly build a GNN-MME describing the full multi-cell system. The FSP of this master equation leads to predicted time-dependent marginal distributions of protein counts in all cells of an arbitrarily connected multi-cell network with arbitrary birth-death parameters.

To overcome this difficulty, we proceed in a different manner. First we use Eq. (2) to derive an exact master equation describing the time-evolution of the marginal distribution *P*(*n_i_, t*) of protein numbers in cell *i* (see Methods)

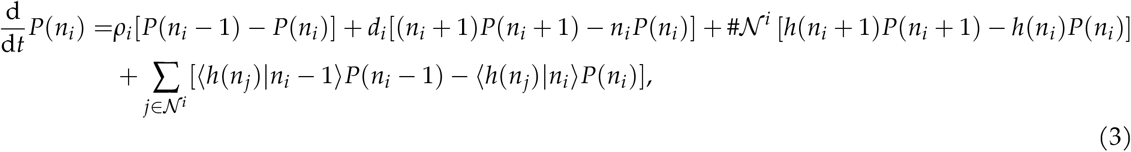

where time *t* is suppressed for convenience, 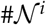 is the number of neighbours of cell *i* and 〈*h*(*n_j_*)|*n_i_*)〉 = ∑*n_j_ h*(*n_j_*)*P*(*n_j_*|*n_i_*) is the the effective transport propensity describing the influx of molecules from cell *j* to cell *i*. Since generally the explicit dependence of the latter on *n* is unknown, it follows that an exact solution of Eq. (3) is impossible. Instead we hypothesize that a neural network’s ability to perform universal function approximation [39] may be conveniently used to establish a mapping from *P*(*n_i_*) and *P*(*n_j_*) to the effective transport propensity 〈*h*(*n_j_*)|*n_i_*〉. This implies that Eq. (3) is now transformed into a set of closed equations for *P*(*n_i_*). It then follows that if we apply the FSP algorithm on a truncated state space, assuming a maximum molecule number *N* – 1, we would need to solve *Nm* coupled differential equations. This is a vast improvement over the *N^m^* equations needed to solve the original stochastic description. This solution is acceptable provided of course we are not interested in the joint distribution function but *only* in the marginals.

Formalising this reasoning, it follows that Eq. (3) can be approximated by a new master equation of the form:

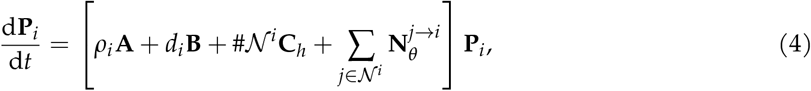

where the vector **P***_i_* collects all the probability elements of cell *i*, i.e., 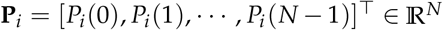 and the matrices are defined as

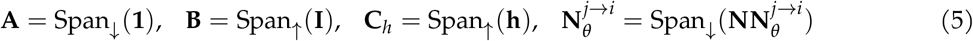

with 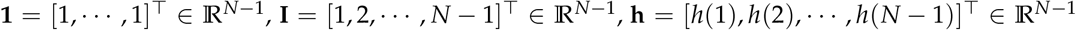 and 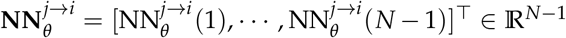 are the neural network outputs. The definitions of the two operators Span_↑_(·) and Span_↓_(·), both of which map a vector **a** = [*a*1,…, *a*_*n*-1_]^⊤^ into sparse matrices, are given by

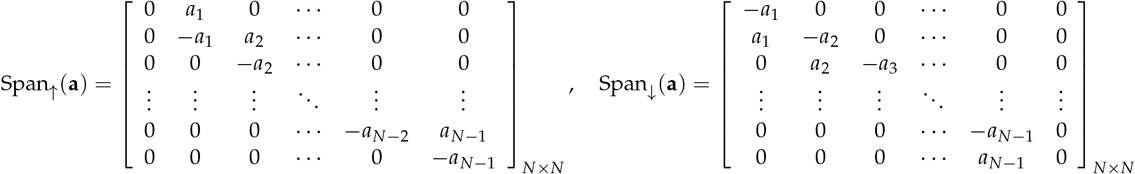

Specifically, the neural network *f_θ_* parametrised by neural network coefficients *θ* (the weights and biases) establishes the mapping

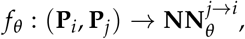

which is independent of the birth-death kinetic parameters *ρ_i_* and *d_i_*. We shall refer to Eq. (4) as the *graph neural network modularised master equation* (*GNN-MME*). See Fig. S1 for an illustration and interpretation of Eq. (4) in the framework of GNNs. We also note that our approach falls under the umbrella of Universal Differential Equations [40] and it is similar to a special type of GNN called a graph network [41].

Finally we devise an effective neural network training procedure. To keep the computational cost to a minimum, we propose to learn the neural network coefficients *θ* of the multi-cell system using SSA simulations of a system with only two neighbouring cells for different sets of birth-death kinetic parameters *ρ_i_* and *d_i_* but with constant transport parameters *D* and *K* (Figs. 1b,c). From these simulations we obtain the time-dependent marginal distributions of protein counts in the two cells, 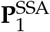 and 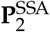. Subsequently, the optimal neural network coefficients are found by minimising a loss function between the latter and the time-dependent distributions given by the FSP solution of the GNN-MME for a two-cell system (Fig. 1d left panel). This completes the learning of the effective propensity. By substituting the latter in a GNN-MME for a multi-cell network we can predict the stochastic dynamics for any arbitrarily connected network (Fig. 1d right panel).

The reliability of this training procedure is tested next. Using three parameter sets (Table S1) we verified that once the neural network was well trained, the FSP solution of the GNN-MME gave time-dependent marginal distributions of protein counts in the two cells that were practically indistinguishable from those obtained using the SSA (Fig. 2a). Subsequently we tested if given the learnt effective propensity (from a two-cell system) we could directly build a GNN-MME to predict the stochastic dynamics of an arbitrarily connected multi-cell network and using birth-death parameters different than the three training sets but with the same transport parameters (Table S2). In Fig. 2c, Figs. S2 and S3 we show the predictions of the GNN-MME for networks of cells connected together in the shape of five letters (Fig. 2b). In all cases, we found that the marginal distribution of protein counts for any cell, at any point in time, agrees very well with distributions obtained from SSA simulations of the multi-cell networks.

**Figure 2:**
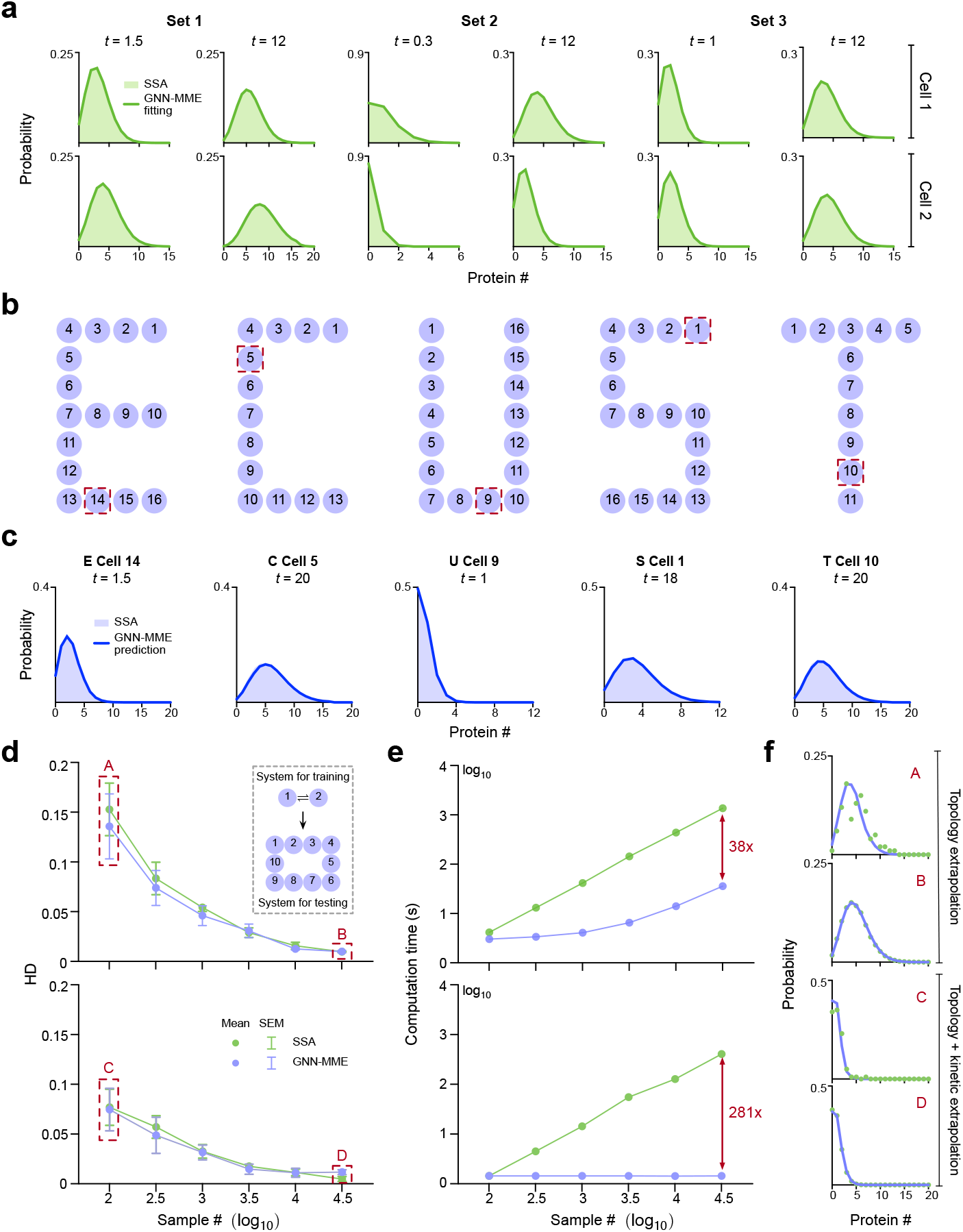
Evaluation of the performance of the GNN-MME for the system of connected cells shown in Fig. 1a. (a) The neural network was trained on two-cell SSA simulations with three different sets of kinetic parameters (Table S1). The optimisation algorithm finds parameters of the neural network that leads to an excellent agreement between the distributions predicted by the two-cell GNN-MME and the SSA distributions used for training. (b,c) The multi-cell GNN-MME with neural networks trained from the 3 parameter sets of two-cell systems accurately predicts the protein distributions for cells in 5 different letter-like topologies where the birth-death kinetic parameters were chosen from uniform distributions 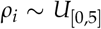 and 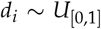 while the transport parameters were fixed to *D* = 10 and *K* = 5 (Table S2). (d-f) (Top panels) We furthermore tested the accuracy and computational performance of the GNN-MME using a circular system of 10 identical cells with *ρ_i_* = 2.5, *d_i_* = 0.5 ∀*_i_*, *D* = 10 and *K* = 5. The GNN-MME trained with a specified number of SSA samples of a two-cell system gave steady-state marginal distributions of comparable accuracy to those directly obtained using the same number of SSA samples of the 10-cell system – the accuracy was measured by the Hellinger distance (HD) between the distributions and the ground truth (a 10-cell SSA simulation with 10^7^ samples). However the computation time of the GNN-MME (time to compute the specified SSA samples for the two-cell system plus the time for the GNN-MME to make predictions) is typically considerably less than that of the same number of 10-cell SSA simulations. (d-f) (Lower panels) Same as for the top panels but where the system for testing had different parameters (*ρ_i_* = *d_i_* = 1 ∀_*i*_) unused for training. Note that in this case the computation time is purely the GNN-MME prediction time since SSA data was not collected. For a fair comparison, benchmarking was performed on one core of an Intel Xeon Gold 6246R 3.40GHz × 64-core CPU 376.6G RAM. See Methods for technical specifications of the neural networks.

We emphasise that because the RDME cannot be easily solved exactly using common analytical methods, the neural network’s success cannot be attributed to a trivial form of the joint distribution *P*(*n*_1_, *n*_2_,…, *n_m_*). The only trivial steady-state solution (product Poisson joint distribution) occurs when the transport propensity *h*(*n*) is linear; but this is not our case since *h*(*n*) is non-linear and in fact the Fano factor (variance divided by the mean) of protein counts in each cell is greater than 1 in steady-state conditions (Figs. S2 and S3).

We have shown that the GNN-MME accurately approximates the marginal protein distribution solutions of the RDME Eq. (3). The remaining question is why our neural-network based approach would be a useful strategy compared to conventional numeric or simulation methods. Earlier we discussed the vast improvement compared to a direct application of the FSP to solve Eq. (3). Now we seek to understand the computational advantage that the GNN-MME approach affords over the direct computation of the distributions of a multi-cell cellular system from a large number of SSA samples. In Figs. 2d-f (top panel), we compare the two methods’ accuracy of the marginal distributions and the computation time for a system of 10 cells connected as a ring and each having the same parameters. The two methods provide comparable accuracy (relative to the “true” solution) given the two-cell SSA simulations used for training the GNN-MME have the same sample size as the 10-cell SSA simulations. However the use of the GNN-MME approach leads to computational savings. Here we note that in our estimation of the computation time for the GNN-MME we did not factor in the training time for neural networks. While this maybe important in other studies, here it is not particularly relevant because once the neural network is trained using two-cell SSA simulations, it can cast predictions on an infinite number of larger scale systems and hence effectively the training time for any given large scale system (such as the 10-cell system in this example) is negligible. In Figs. 2d-f (bottom panel), we repeat the comparison of the accuracy and the computation time but now for the more challenging case where the parameters of the GNN-MME are different than those used for the SSA training of the two-cell system. Use of the GNN-MME approach leads to an impressive ~ 300 times decrease in the computation time compared to the SSA of the full system while maintaining similar accuracy.

### The neural network based approach is extensible to multi-species systems: spatio-temporal mRNA & protein patterns at subcellular resolution

Thus far we have illustrated the GNN based approach using a single-species system. We next consider its generalisation to a multi-species system which will require the use of additional neural networks. To be specific, consider a spatial model of a genetic negative feedback loop. We divide (the 2D cross section of) a cell into a system of 11 × 11 voxels; a voxel represents either a small volume around the gene locus, or a small spatial domain in the nucleus or cytoplasm. We define a set of reactions to model the following phenomena. Messenger RNA (mRNA) is produced at the gene locus with a rate that is inversely proportional to the number of proteins in the vicinity of the locus, mRNA diffuses in the nucleus and can be transported into the cytoplasm where it is translated into protein and it can also degrade. Proteins diffuse in the cytoplasm and can also be transported to the nucleus where they can diffuse to the gene locus and affect the mRNA production rate. Note that by diffusion here we mean simple diffusion, not the active transport we considered earlier. This system of reactions constitutes a negative feedback loop since an increase in the mRNA levels leads to a similar increase in protein levels which after a time lag leads to a decrease in the mRNA levels. The fine-grained cell and the associated reactions are illustrated in Fig. 3a.

**Figure 3:**
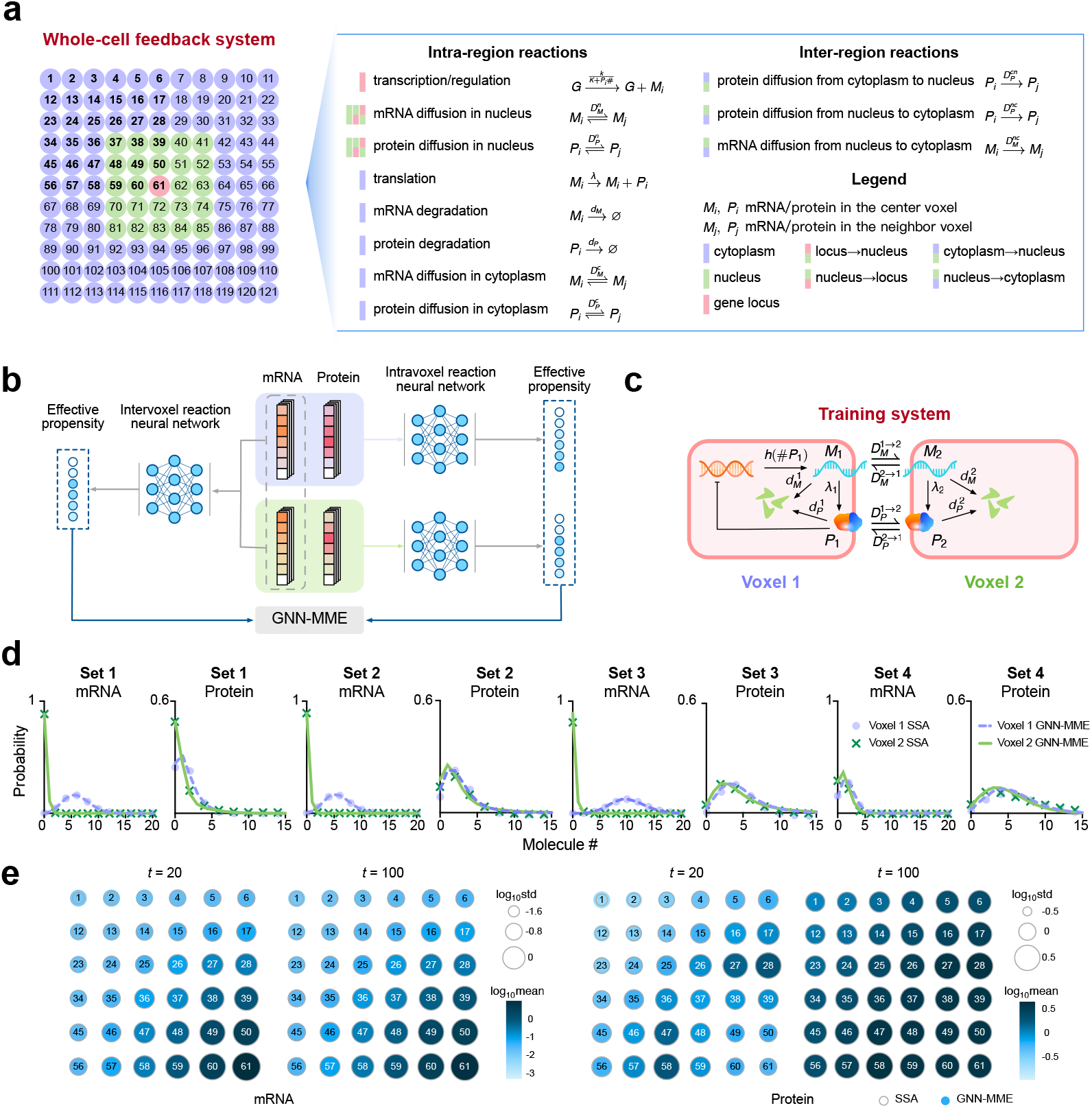
GNN-MME approach enables accurate prediction at sub-cellular resolution of mRNA and protein dynamics across a whole cell from stochastic simulations of a two-voxel system. (a) The 2D cross section of a cell is divided into 11 × 11 voxels; the central 5 × 5 voxels represent the nucleus (green), the center represents a small nuclear region containing the gene locus (red), and the rest represent the cytoplasm (purple). Reactions are defined describing transcriptional regulation via protein binding, translation, diffusion and degradation. mRNA degradation can only occur in the cytoplasm and while protein can move in and out of the nucleus, mRNA movement is only allowed from the nucleus to the cytoplasm. (b) We use neural networks to learn an approximate mapping (i) from the marginal distributions of mRNA or protein in two adjacent voxels (shown in blue and green) to the effective influx diffusion propensity (only one of these two intervoxel reaction neural networks is shown) and (ii) from the marginal distributions of mRNA and protein in the same voxel to the effective transcription and translation propensities. These propensities enable us to construct a GNN-MME (Eqs. (6) and (7)) whose solution using FSP directly gives approximate marginal distributions of mRNA and protein. (c) Training of the neural networks is performed using a two-voxel system which while much simpler than the whole cell system, it describes the same phenomena but on a much smaller spatial scale. (d) SSA simulations of the two-voxel system with four sets of parameters (Table S3) were used to train the intra- and intervoxel neural networks. The optimisation algorithm finds parameters of the neural networks that lead to excellent agreement between the distributions predicted by the two-voxel GNN-MME (Eq. (10)) and the SSA distributions used for training. (e) The propensities learnt from the two-voxel simulations are used to construct the GNN-MME of the whole cell system in (a) (parameters in Table S4). Its FSP solution leads to time-dependent protein and mRNA distributions whose first and second moments are in good agreement with SSA (10^5^ samples) of the whole cell system - the color and size of the shaded circles show the logarithmic mean and the logarithmic standard deviation (std) predicted by the GNN-MME, respectively; the open grey circles show the logarithmic std predicted by SSA. For a comparison of the distributions see Figs. S4-S6. See Methods for technical specifications of the neural networks.

If we use the same GNN approach as in the previous single-species example, the resulting GNN-MME is a set of equations for the joint probability distribution of mRNA and protein numbers in each voxel. This implies we need to solve 121*N*^2^ differential equations simultaneously, which while it is a significant improvement over the *N*^242^ differential equations solved using the standard FSP, nevertheless it is still quite a large number if *N* is larger than a handful of molecules. To further reduce the computation time, what we need are differential equations for the marginal distributions of each of the two species in each voxel. To achieve this aim, we first write the exact master equation for the marginal distributions of mRNA and protein in each voxel and then construct a GNN-MME by means of two neural networks: (i) an intervoxel reaction neural network is used to find an approximate mapping from the marginal distributions of mRNA or protein in the two adjacent voxels to the effective influx diffusion propensity; (ii) an intravoxel reaction neural network is used to find an approximate mapping of the marginal distributions of mRNA and protein in the same voxel to the effective transcription and translation propensities. The construction of the GNN-MME is illustrated in Fig. 3b. Specifically, for voxels in the nucleus (including the gene locus), the GNN-MME takes the form

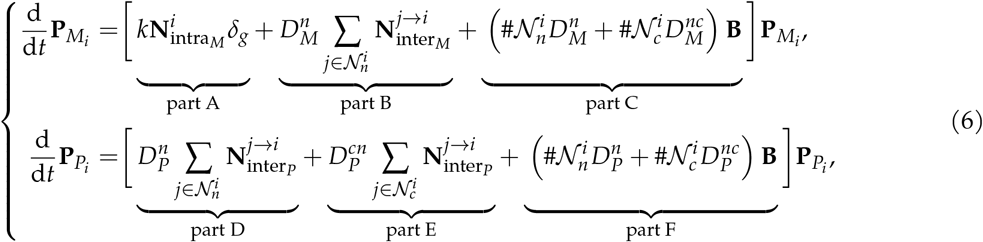

while in the cytoplasm it has the form

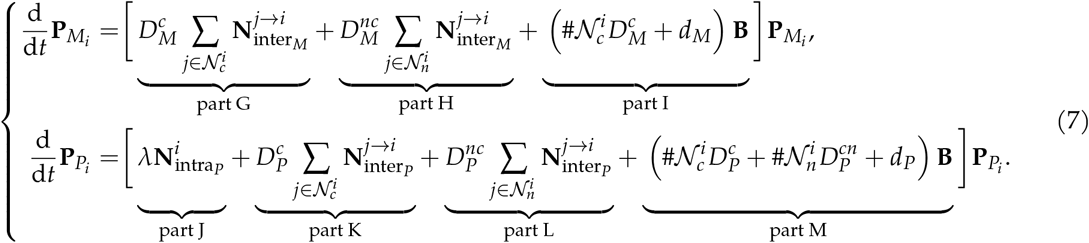

Here **P***_M_i__* and **P***_P_i__* denote the marginal distributions of mRNA and protein counts in voxel *i*, respectively. The reactions associated with the various rate parameters in these equations can be found in Fig. 3a. The symbols 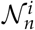 and 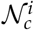 mean the set of neighbours in the nucleus and cytoplasm for voxel *i* respectively. In Eq. (6), part A uses an intravoxel neural network to model the effect of protein-modulated transcription at the gene locus voxel (*δ_g_* is a delta function and equal to 1 if and only if voxel *i* is the gene locus), parts B and D use two intervoxel neural networks to describe the net diffusive influx of mRNA and protein into a nuclear voxel *i* respectively, parts C and F describe the net diffusive outflux of mRNA and protein from a nuclear voxel *i* respectively, and part E uses an intercellular neural network to describe the diffusive influx of protein from adjacent cytoplasmic voxels to the nuclear voxel *i*. In Eq. (7), parts G and H describe the diffusive influx of mRNA from adjacent cytoplasmic and nuclear voxels to cytoplasmic voxel *i* respectively, the same for parts K and L but for protein, part J uses an intravoxel network to describe translation, part I describes diffusive outflux and degradation of mRNA in a cytoplasmic voxel, and part M describes the same effects but for protein.

The matrix **B** is the same as defined in Eq. (5). The four neural network matrices in Eqs. (6) and (7) are 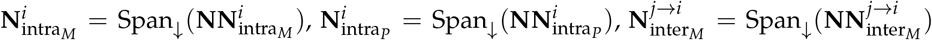 and 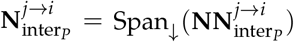. The associated four neural networks learn the following mappings

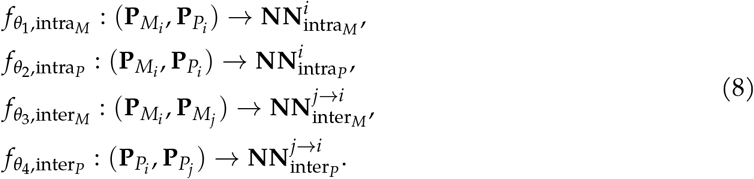

We emphasise that given Eqs. (6) and (7), the number of differential equations that need to be solved is reduced from *N*^242^ (standard FSP) to just 242*N*.

What remains is to define a training protocol so that we can efficiently learn the optimal neural network coefficients *θ* = [*θ*_1_, *θ*_2_, *θ*_3_, *θ*_4_]*^⊤^* from stochastic simulations of a system that is much smaller in scale than the whole cell system but which effectively models the same phenomena. Thus for training purposes, we consider a two-voxel, coarse-grained version of the whole cell model with the following set of reactions (also illustrated by a cartoon in Fig. 3c).

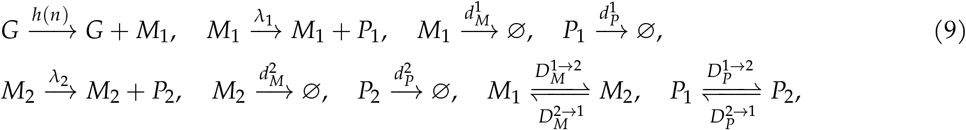

where 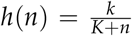. To train the four neural networks in (8), we need to derive the GNN-MME for the two-voxel system, which is indeed a reduced version of Eqs. (6) and (7):

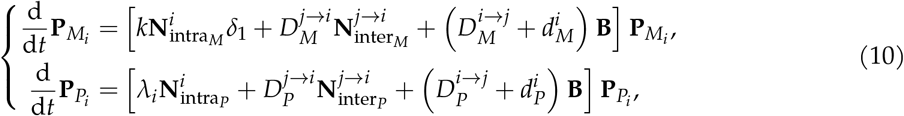

for *i* = 1, 2 and 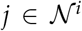. The delta function *δ*_1_ = 1 if and only if *i* = 1. The training dataset consists of the SSA marginal distributions of mRNA and protein for both voxels for four different sets of kinetic parameters (Table S3). In Fig. 3d we confirm that the neural networks are well trained by an excellent agreement between the GNN-MME for the two-voxel system and the SSA distributions used for training.

Using the learnt propensities from the two-voxel system, we then build the GNN-MME for the whole cell 121-voxel system shown in Fig. 3a, i.e. Eqs. (6)-(7). Note that some of the parameters of the whole cell system (Table S4) are different than those used for training the two-voxel system. In Fig. 3e we show the mean and variance of mRNA and protein counts for a quadrant of the whole cell system (the other quadrants show the same information because of the spatial symmetry of the system) at two different times; for the corresponding marginal distributions see Figs. S4-S6. The excellent agreement with the distributions computed using the SSA of the whole cell system confirms the GNN-MME’s ability to accurately predict the fine-grained spatio-temporal pattern of mRNA and protein dynamics at sub-cellular resolution from SSA stochastic trajectories of a coarse-grained, two-voxel model. The results also identify an interesting dependence of the Fano factor of protein fluctuations with distance from the gene locus, e.g. in Fig. S5 from voxel 56 to 61, the Fano factor increases as we move from the cell boundary to the nucleus-cytoplasm boundary then decreases as we move into the nucleus and finally again increases as it approaches the gene locus.

## Discussion

In this article, we have described a novel method which uses GNNs to automatically “derive” a master equation describing the time-evolution of the marginal probability distributions of gene products in each voxel of a spatially extended system. The numerical solution of this effective master equation leads to solutions for the marginal distributions in a small fraction of the time of conventional methods, thereby potentially enabling the large scale simulation of stochastic gene expression at the sub-cellular resolution for every cell in a tissue.

While it is possible to write a master equation for the marginal distributions from first principles, this equation cannot be generally solved because it is not closed due to the presence of terms which are not functions of the marginal distributions. We circumvent this issue by using a neural network to learn (from a small batch of stochastic simulations) these terms as functions of the marginal distributions, thereby closing the master equation and enabling its solution using FSP. Whilst we showcased the procedure using one- and two-species examples, it is general for a system with any number of voxels and species. It is straightforward to deduce that the direct solution of this effective master equation for a spatially extended system with *M* voxels and *S* species, assuming an upper bound *N* – 1 for the molecule counts in each voxel, implies the solution of *NMS* coupled ordinary differential equations. By contrast, solving the reaction-diffusion master equation describing the full spatial stochastic dynamics using FSP would imply solving *N^MS^* ordinary differential equations. To put in perspective, if we want to model the spatial variation of transcript numbers of one gene inside a single cell which has been discretised into 100 voxels, with each voxel holding at most 1 transcript, the conventional FSP requires the solution of 10^30^ coupled differential equations while our method reduces the computation to the solution of 200 coupled differential equations! We also showed that the method’s computation time is often much smaller than Monte Carlo methods such as the SSA - fundamentally this is because the neural network only needs two-voxel simulations to train it and then it can extrapolate the predictions to an arbitrarily connected voxel network with different kinetic parameter values. The method is promising for whole cell modelling since we showed that neural network training on a coarse-grained, two-voxel version of a cell is sufficient to cast accurate predictions for much finer discretisations of intracellular space.

A limitation of our method is that it cannot give the joint distributions of molecular counts in all voxels. The method can either give the marginal distributions of each species in each voxel or else if one use the neural network to only approximate the transport influx terms then the method gives the joint distributions of all species in each voxel. We do not see this as a major hindrance since often this is all the information that is needed. We note that while it might be possible for some systems to analytically approximate the terms in the master equations for the marginals [42–44] which the GNN targets, such a procedure would be based on some implicit assumption (akin to moment-closure techniques) and hence its accuracy would generally be less compared to the automated technique we have described.

Recently a range of neural network based methods have been devised which also significantly reduce the computational burden compared to conventional simulation methods [45–49]. A feedforward neural network was used in [45] to construct Markovian approximations of stochastic biochemical reaction systems with non-Markovian dynamics; this method is also FSP-based as the method developed in this paper. DeepCME [46] uses reinforcement learning to estimate the moments of molecular counts from stochastic simulations. Mixture Density Networks have been used to learn the transition kernel of the chemical master equation [47]. Nessie (Neural Estimation of Stochastic Simulations for Inference and Exploration) [48] uses a neural-network approach to learn a negative binomial mixture approximation to the marginal distributions of each species. A similar approach to the latter but extending the prediction to steady-state joint distribution solutions for a two-species model of the life cycle of RNA was recently reported in [49]. We emphasise that in all these cases, the systems of interest are non-spatial (well-mixed) chemical reaction systems. While in principle these methods could also be applied to approximating spatially-extended systems, in practice this would be considerably difficult because the neural networks in these methods try to learn a mapping from the very large dimensional parameter space (which scales with the number of voxels) to the joint or marginal distributions (or their moments). By contrast, our method bypasses this difficulty by training a neural network using two-voxel simulations to “derive” a master equation for the marginal distributions in each voxel of an arbitrarily large and complex spatially-extended system. Finally we note that by optimising not only the weights and biases of the neural networks but also simultaneously for the kinetic parameter values, it might be possible to use our method together with spatial genomics data to learn cell-specific transcriptional and transport parameters.

## Methods

### Derivation of the marginal master equation Eq. (3)

By taking the sum of *n_k_, k* ≠ *i* on both sides of Eq. (2), we obtain

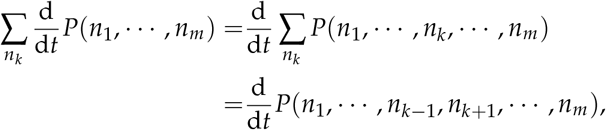

where the last step follows from the law of total probability. By sequentially repeating the above procedure for all *n_k_, k* ≠ *i*, the term becomes

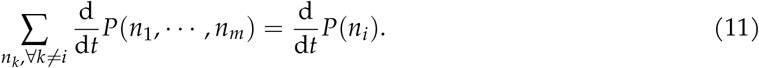

For the first term on the right hand side of Eq. (2), taking the sum over *n_k_* and using the same procedure, we obtain

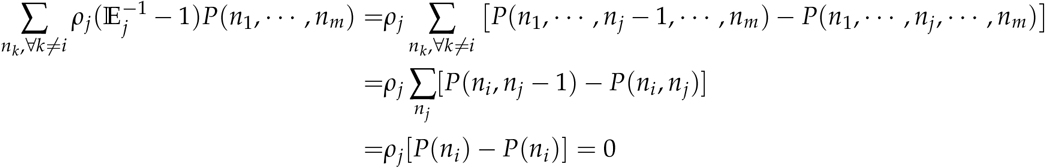

for *j* ≠ *i*, and

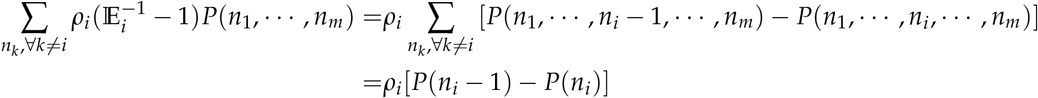

for *j = i*. These results imply

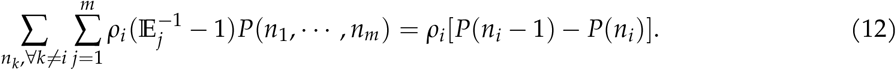

Similarly, we have

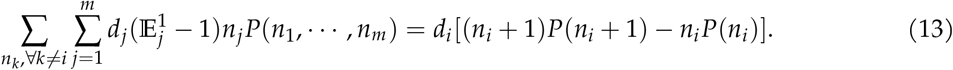

Subsequently, it follows that for *ℓ* ≠ *i* and 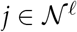

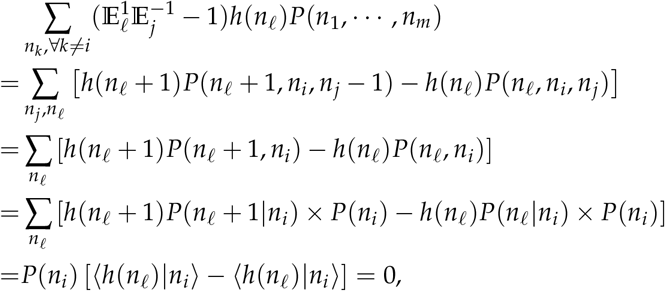

and for *ℓ* = *i* and 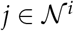

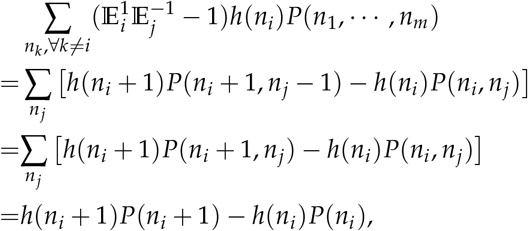

both of which imply that

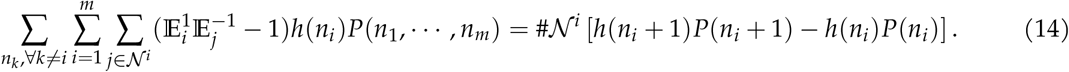

Lastly, we perform marginalisation of the last term on the right hand side of Eq. (2). It follows that for *ℓ* = *i* and 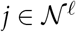

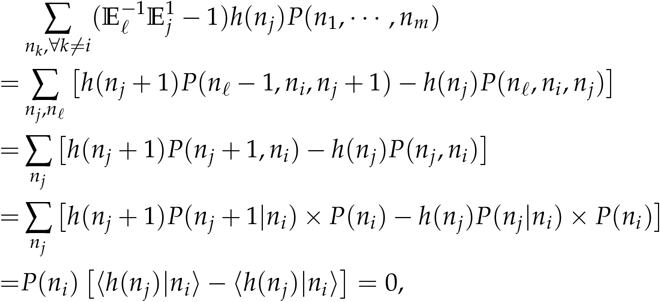

and for *ℓ* ≠ *i* and 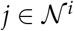,

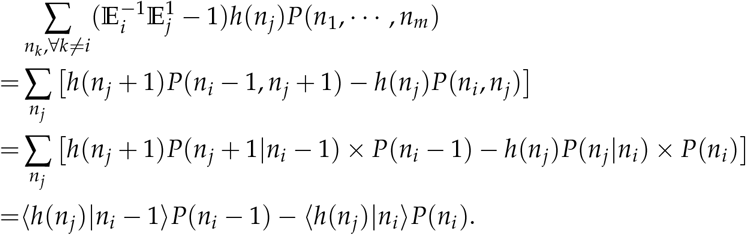

These imply that

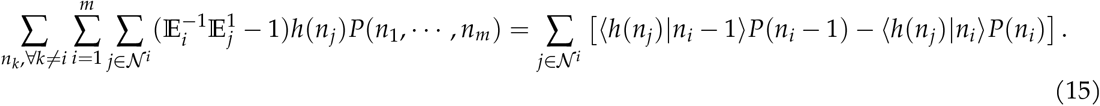

### Neural network specifications

In the example of birth-death reactions in a system of connected cells, the truncation number for the protein count distribution is *N* = 39. The neural network takes the marginal distributions of two adjacent cells as inputs and outputs the effective transport influx propensity into each voxel. The neural network has 3 layers with 80-20-39 neurons from input to output respectively, while the activation functions are `tanh` and `LeakyReLU` accordingly. The two-cell distribution snapshots used for training are evenly distributed over time *t* = 20 with time interval *t_s_* = 0.1. The SSA sample number is 8 × 10^4^ for Figs. 2a-c. The initial condition is zero protein in all cells. The loss function is the sum of the the mean squared difference between the distributions predicted by the FSP of the two-voxel GNN-MME and the SSA distribution snapshots; this sum is over all time points, kinetic parameter sets and cells. The minimisation of the loss function is performed by gradient based optimisation algorithms. Specifically, gradients are computed by automatic differentiation via packages `DiffEqSensitivity` and `Zygote`, backpropagation is used to efficiently compute the gradient of the loss function with respect to the parameters of the neural network and the `Adam` optimiser is then used to update the parameters to minimise loss. The learning rate of the optimiser is set to *η* = 0.01 for 300 epochs.

In the example of whole cell model of a genetic negative feedback loop, the truncation number for the protein count distribution is *N* = 39. For the four neural networks in Eq. (8), *f*_*θ*_1_,*intra_M_*_ and *f*_*θ*_2_,*intra_P_*_ take the marginal distributions of mRNA and protein from the same voxel as inputs and output the effective transcriptional and translational propensities respectively, while *f*_*θ*_3_,*inter_M_*_ and *f*_*θ*_4_,*inter_M_*_ take the marginal distributions of mRNA or protein of two adjacent voxels as inputs and output the effective diffusive influx propensities from the neighbouring cell. The four neural networks have 3 layers with 80-20-39 neurons from input to output respectively, while the activation functions are tanh and `LeakyReLU` accordingly. The two-voxel distribution snapshots are evenly distributed over time *t* = 60 with time interval *t_s_* = 0.5, and the SSA sample number is 8 × 10^4^. The initial condition is zero mRNA and protein. The loss function is similar to what was defined for the previous system. The training is performed by using the Adam optimiser with two stages: the first stage uses a learning rate *η* = 0.01 for 300 epochs, while the second stage uses *η* = 0.001 for another 300 epochs.

## Code availability

The codes, readme file and data for GNN-MME can be found at 10.5281/zenodo.7678839. The codes are implemented by Julia 1.6.5 and its package Flux v0.12.8, `DifferentialEquations` v7.0.0, `DiffEqSensitivity` v6.66.0 and `Zygote` v0.6.33.

## Acknowledgements

We acknowledge Rui Wang for a preliminary study exploring the viability of GNNs in solving stochastic gene expression problems. We thank Kaan Öcal and Augustinas Sukys for comments on the manuscript. W.Z. acknowledges support from the Natural Science Foundation of China (NSFC No. 61925305). Z.C. acknowledges support from NSFC No. 62073137, Shanghai Action Plan for Technological Innovation Grant (No. 22ZR1415300, 22511104000) and Shanghai Center of Biomedicine Development. R.G. acknowledges support from the Leverhulme Trust grant (RPG-2020-327).

## Author contributions

Z.C. and R.G. designed and supervised research, acquired funding and wrote the manuscript with input from the others. R.C., L.X., X.Z., X.F., W.Z. and Z.C. performed research and analysed the data.

---

**Figure S1:**
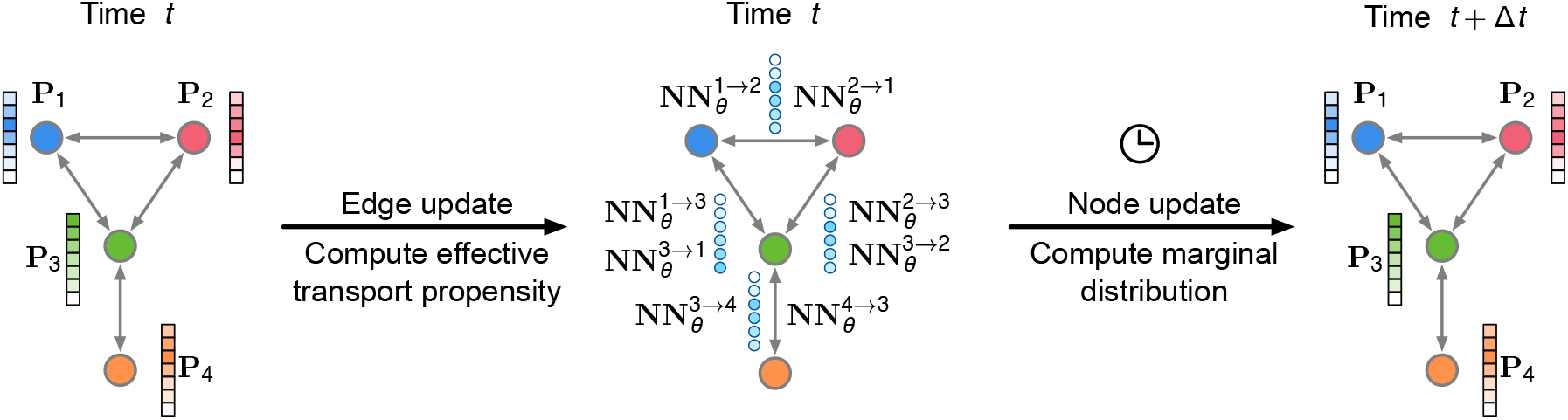
Interpretation of Eq. (4) in the framework of GNN. The physical adjacency among cells/voxels naturally defines the topology (connectivity between cells/voxels), and the neighbour set 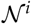 of each cell/voxel mathematically represents the topological encoding. In a graph, there are two components; one is node (a cell or voxel), and the other is edge (the diffusion influx between cells/voxels). Either node or edge has attributes. The node attribute here is the marginal distribution **P**_*i*_ in Eq. (4), while the edge attribute is the effective transport propensity 〈*h*(*n_j_*)|*n_i_*〉 approximated by a multilayer perceptron (MLP) (*f_θ_*), which outputs 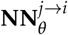. In each time step, edge update is first performed, wherein a MLP takes marginal distributions of two adjacent cells/voxels as input and outputs the effective transport propensity 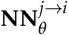. The yielded effective transport propensity (at time *t*) integrated with the marginal master equations Eq. (4) leads to the marginal distribution of each cell/voxel at time *t* + Δ*t*. These two steps can be iterated from time *t* = 0 to any time *T*. Though we use MLPs for approximation, their integration with topology encoding (via the sum over the numbers of neighbouring cells/voxels) naturally makes Eq. (4) a GNN.

**Figure S2:**
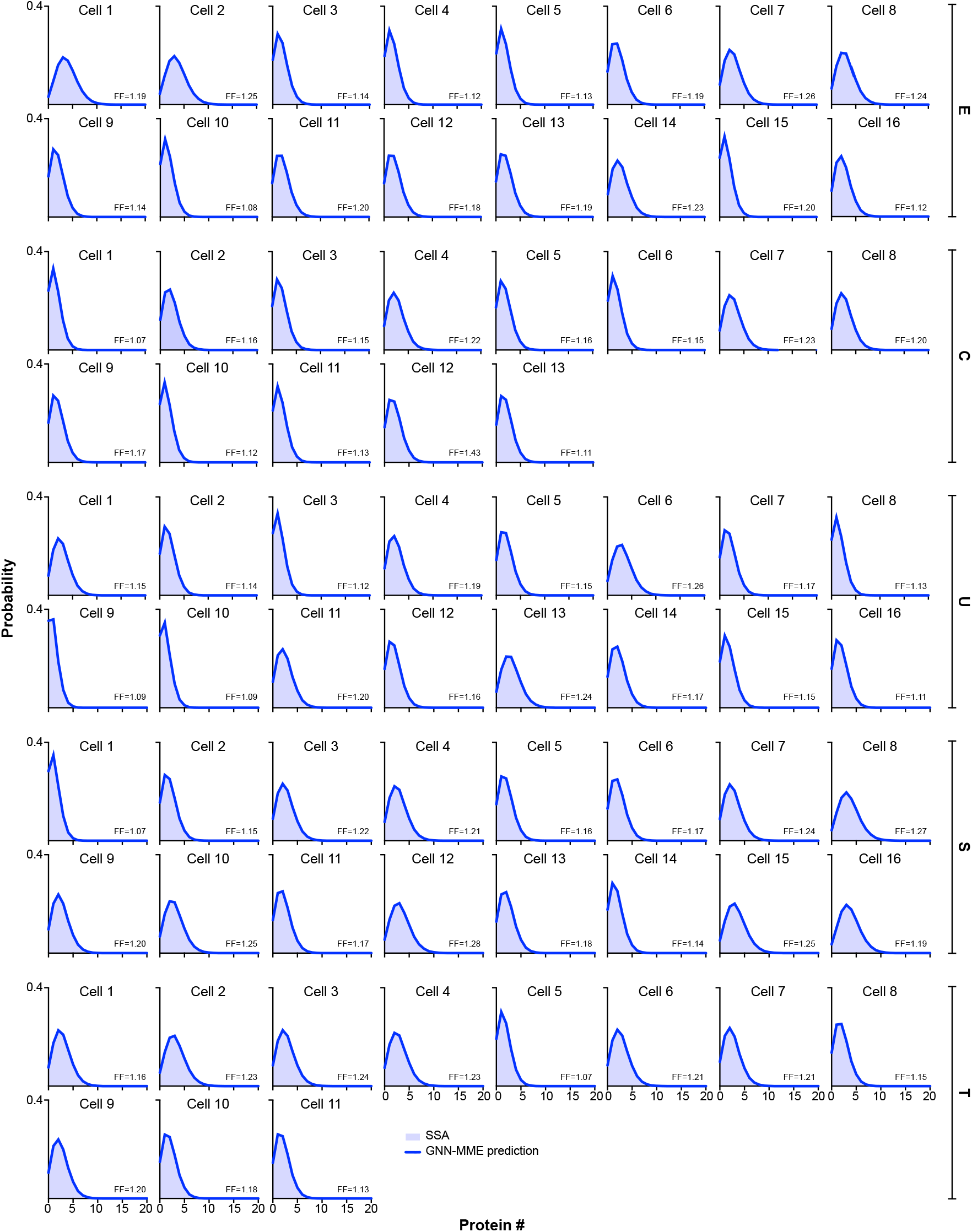
Marginal distributions of protein counts at time *t* = 1.5 for the five letter-like cell networks in Fig. 2b with kinetic parameters given in Table S2. The two-cell GNN-MME was trained using 3 parameter sets given in Table S1. The learnt propensities were then used to build the multi-cell GNN-MME Eq. 4 whose FSP solution accurately matches the SSA distributions for all cells in the five cell network topologies. Note that the Fano factor (FF) of protein fluctuations in each cell is greater than 1. This deviation from Poisson statistics is due to the nonlinearity of the transport propensity function *h*(*n*) = *Dn*/(*K* + *N*).

**Figure S3:**
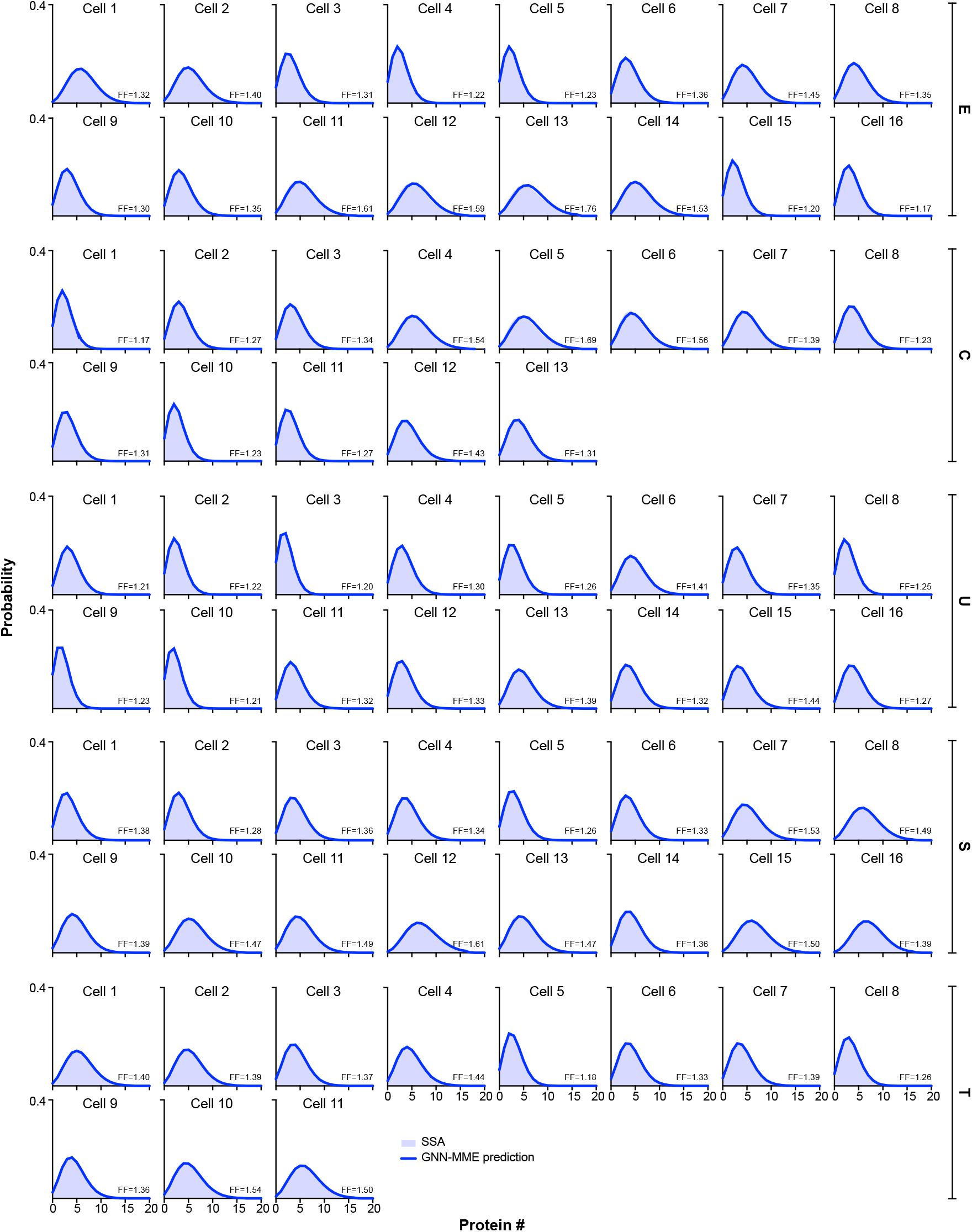
Marginal distributions of protein counts at time *t* = 20 for the five letter-like cell networks in Fig. 2b. All details are the same as in Fig. S2. Note that at *t* = 20 the system has reached steady-state.

**Figure S4:**
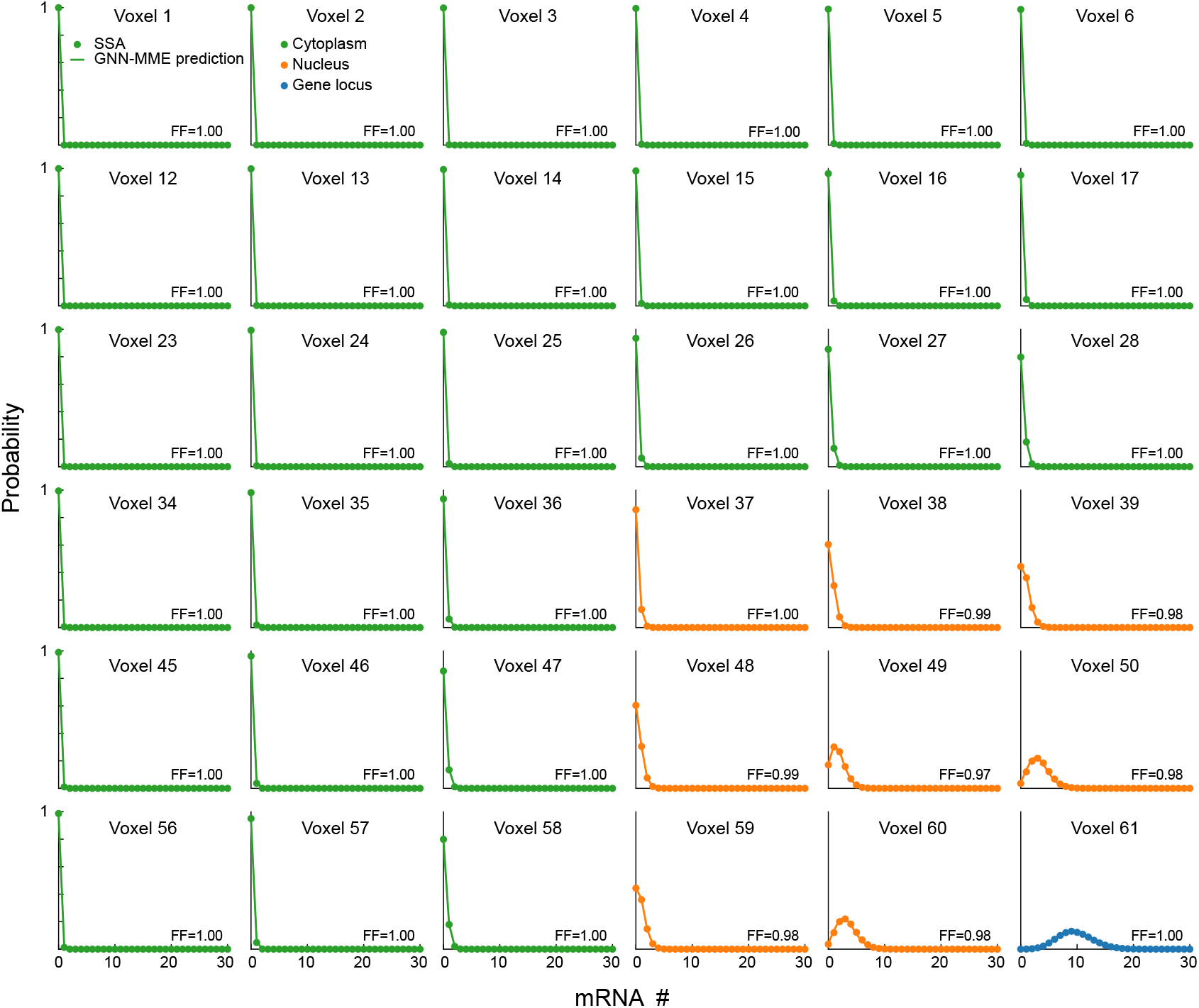
Marginal mRNA distributions at *t* = 20 in each voxel in a quadrant of the whole cell system shown in Fig. 3a with parameters given in Table S4. The results for later times are very similar and hence not shown. The two-voxel GNN-MME Eq. 10 for the system shown in Fig. 3c was trained using 4 parameter sets given in Table S3. The learnt propensities were used to build the whole cell GNN-MME Eq. 6-7 whose FSP solution accurately matches the SSA predictions of the whole cell system. Note that the fluctuations are Poissonian (FF = 1) in the cytoplasm and weakly sub-Poissonian (FF < 1) in the nucleus due to suppression induced by negative feedback.

**Figure S5:**
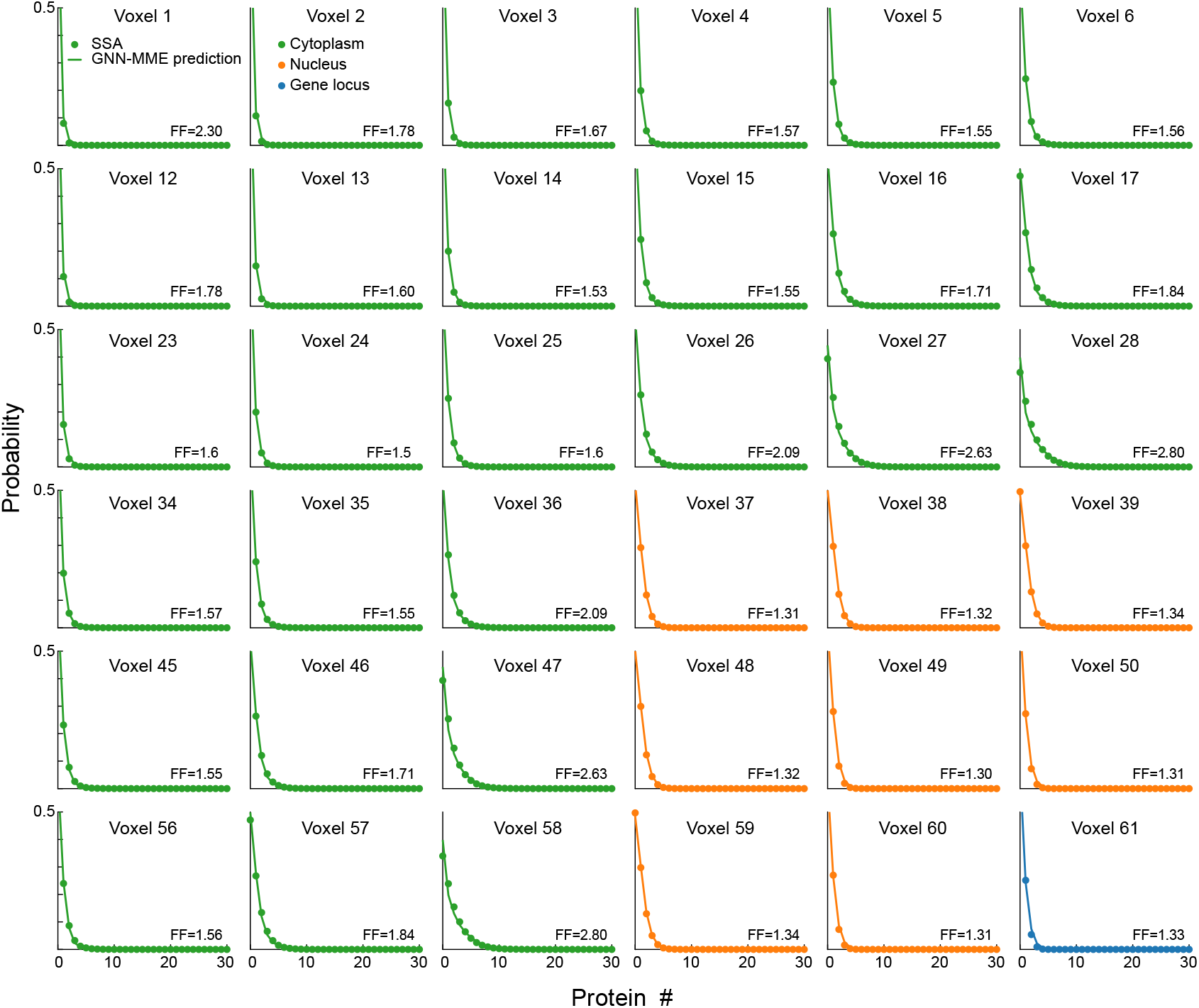
Marginal protein distributions at *t* = 20 in each voxel in a quadrant of the whole cell system shown in Fig. 3a with parameters given in Table S4. Details are the same as in Fig. S4. Note that the fluctuations are super-Poissonian (FF > 1) across the whole cell but the deviations from Poisson behavior are most pronounced in the cytoplasm.

**Figure S6:**
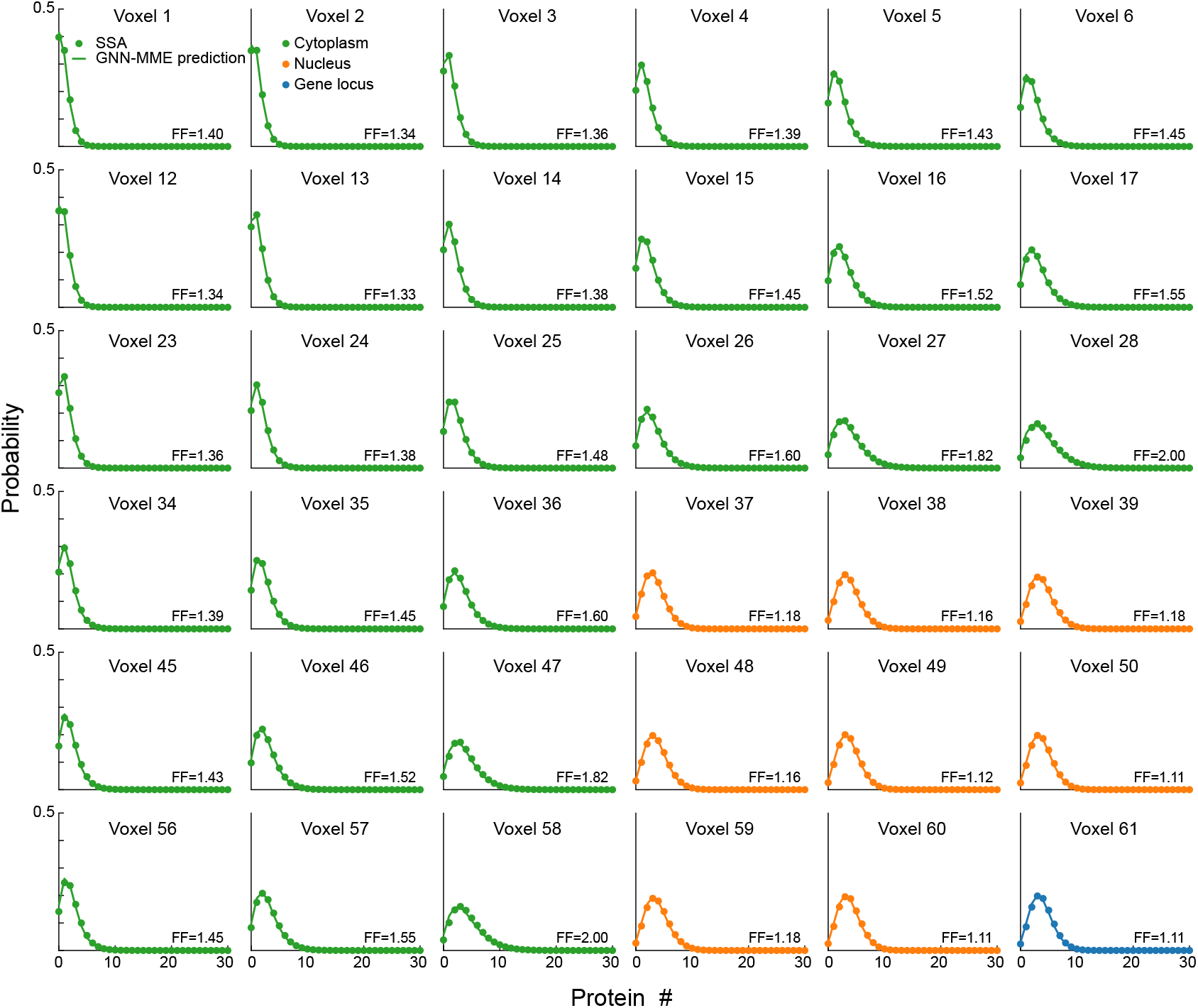
Marginal protein distributions at *t* = 100 in each voxel in a quadrant of the whole cell system shown in Fig. 3a with parameters given in Table S4. Details are the same as in Fig. S4.

**Table S1:** Kinetic parameters of the two-cell SSA simulations used to train the GNN-MME in Fig. 2a

| Datasets | Kinetic parameters |  |  |  |  |  |
| --- | --- | --- | --- | --- | --- | --- |
| | $\rho_1$ | $\rho_2$ | $d_1$ | $d_2$ | $D$ | $K$ |
| Set 1 | 2.616 | 4.841 | 0.614 | 0.443 | 10 | 5 |
| Set 2 | 4.111 | 0.031 | 0.532 | 0.793 | 10 | 5 |
| Set 3 | 2.982 | 3.761 | 0.862 | 0.748 | 10 | 5 |

**Table S2:**
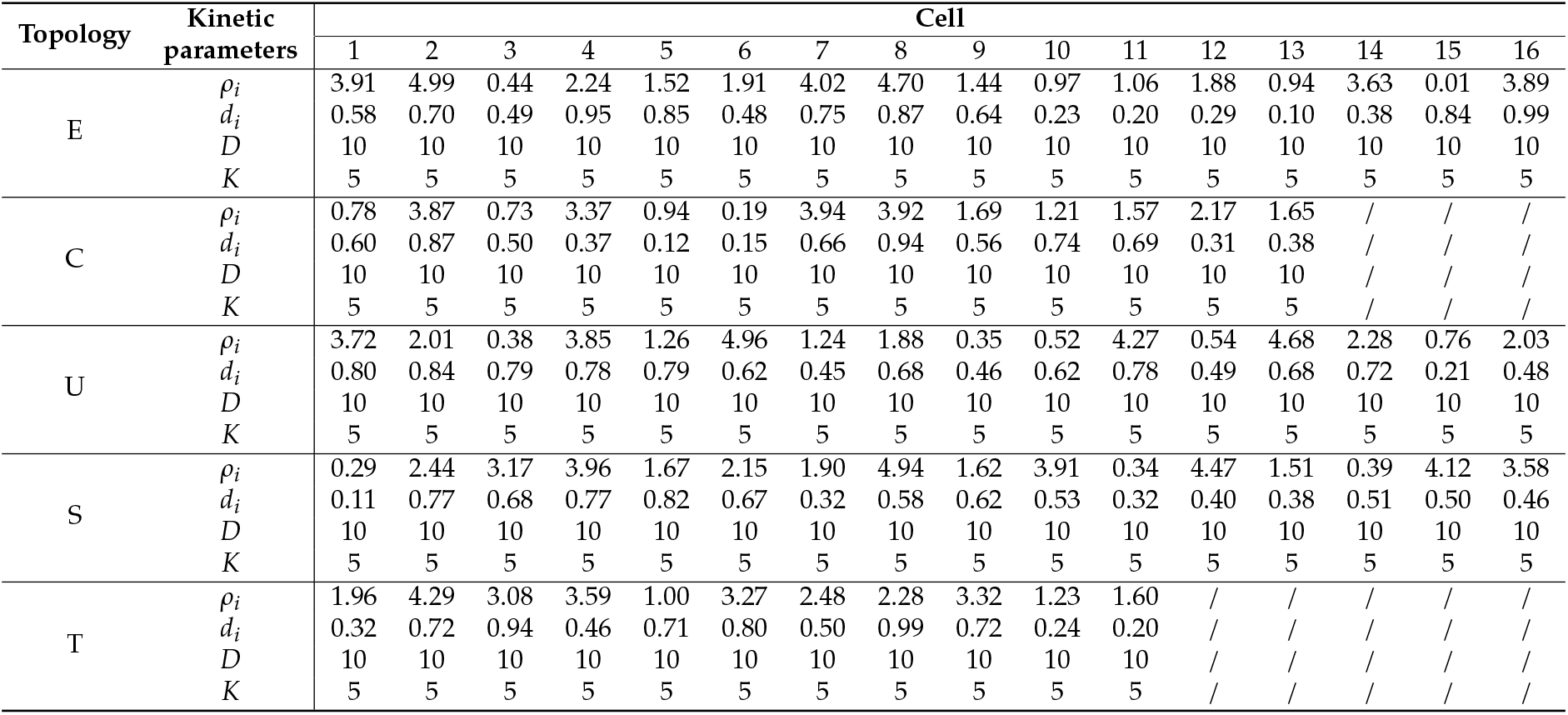
Cellular kinetic parameters of the letter-like cell systems in Fig. 2b.

| Topology | Kinetic parameters | Cell |  |  |  |  |  |  |  |  |  |  |  |  |  |  |  |
| --- | --- | --- | --- | --- | --- | --- | --- | --- | --- | --- | --- | --- | --- | --- | --- | --- | --- |
|  |  | 1 | 2 | 3 | 4 | 5 | 6 | 7 | 8 | 9 | 10 | 11 | 12 | 13 | 14 | 15 | 16 |
| E | $\rho_i$ | 3.91 | 4.99 | 0.44 | 2.24 | 1.52 | 1.91 | 4.02 | 4.70 | 1.44 | 0.97 | 1.06 | 1.88 | 0.94 | 3.63 | 0.01 | 3.89 |
| | $d_i$ | 0.58 | 0.70 | 0.49 | 0.95 | 0.85 | 0.48 | 0.75 | 0.87 | 0.64 | 0.23 | 0.20 | 0.29 | 0.10 | 0.38 | 0.84 | 0.99 |
| | $D$ | 10 | 10 | 10 | 10 | 10 | 10 | 10 | 10 | 10 | 10 | 10 | 10 | 10 | 10 | 10 | 10 |
| | $K$ | 5 | 5 | 5 | 5 | 5 | 5 | 5 | 5 | 5 | 5 | 5 | 5 | 5 | 5 | 5 | 5 |
| C | $\rho_i$ | 0.78 | 3.87 | 0.73 | 3.37 | 0.94 | 0.19 | 3.94 | 3.92 | 1.69 | 1.21 | 1.57 | 2.17 | 1.65 | / | / | / |
| | $d_i$ | 0.60 | 0.87 | 0.50 | 0.37 | 0.12 | 0.15 | 0.66 | 0.94 | 0.56 | 0.74 | 0.69 | 0.31 | 0.38 | / | / | / |
| | $D$ | 10 | 10 | 10 | 10 | 10 | 10 | 10 | 10 | 10 | 10 | 10 | 10 | 10 | / | / | / |
| | $K$ | 5 | 5 | 5 | 5 | 5 | 5 | 5 | 5 | 5 | 5 | 5 | 5 | 5 | / | / | / |
| U | $\rho_i$ | 3.72 | 2.01 | 0.38 | 3.85 | 1.26 | 4.96 | 1.24 | 1.88 | 0.35 | 0.52 | 4.27 | 0.54 | 4.68 | 2.28 | 0.76 | 2.03 |
| | $d_i$ | 0.80 | 0.84 | 0.79 | 0.78 | 0.79 | 0.62 | 0.45 | 0.68 | 0.46 | 0.62 | 0.78 | 0.49 | 0.68 | 0.72 | 0.21 | 0.48 |
| | $D$ | 10 | 10 | 10 | 10 | 10 | 10 | 10 | 10 | 10 | 10 | 10 | 10 | 10 | 10 | 10 | 10 |
| | $K$ | 5 | 5 | 5 | 5 | 5 | 5 | 5 | 5 | 5 | 5 | 5 | 5 | 5 | 5 | 5 | 5 |
| S | $\rho_i$ | 0.29 | 2.44 | 3.17 | 3.96 | 1.67 | 2.15 | 1.90 | 4.94 | 1.62 | 3.91 | 0.34 | 4.47 | 1.51 | 0.39 | 4.12 | 3.58 |
| | $d_i$ | 0.11 | 0.77 | 0.68 | 0.77 | 0.82 | 0.67 | 0.32 | 0.58 | 0.62 | 0.53 | 0.32 | 0.40 | 0.38 | 0.51 | 0.50 | 0.46 |
| | $D$ | 10 | 10 | 10 | 10 | 10 | 10 | 10 | 10 | 10 | 10 | 10 | 10 | 10 | 10 | 10 | 10 |
| | $K$ | 5 | 5 | 5 | 5 | 5 | 5 | 5 | 5 | 5 | 5 | 5 | 5 | 5 | 5 | 5 | 5 |
| T | $\rho_i$ | 1.96 | 4.29 | 3.08 | 3.59 | 1.00 | 3.27 | 2.48 | 2.28 | 3.32 | 1.23 | 1.60 | / | / | / | / | / |
| | $d_i$ | 0.32 | 0.72 | 0.94 | 0.46 | 0.71 | 0.80 | 0.50 | 0.99 | 0.72 | 0.24 | 0.20 | / | / | / | / | / |
| | $D$ | 10 | 10 | 10 | 10 | 10 | 10 | 10 | 10 | 10 | 10 | 10 | / | / | / | / | / |
| | $K$ | 5 | 5 | 5 | 5 | 5 | 5 | 5 | 5 | 5 | 5 | 5 | / | / | / | / | / |

**Table S3:** Kinetic parameters for SSA simulations training the GNN-MME of the two-voxel system in Figs. 3c,d

|  | Intracellular kinetic parameters |  |  |  |  |  |  |  | Intercellular kinetic parameters |  |  |  |
| --- | --- | --- | --- | --- | --- | --- | --- | --- | --- | --- | --- | --- |
| | $k$ | $K$ | $\lambda_1$ | $\lambda_2$ | $d_M^1$ | $d_M^2$ | $d_P^1$ | $d_P^2$ | $D_M^{1 \rightarrow 2}$ | $D_M^{2 \rightarrow 1}$ | $D_P^{1 \rightarrow 2}$ | $D_P^{2 \rightarrow 1}$ |
| Set 1 | 40 | 15 | 0.1 | 4 | 0.4 | 0.2 | 0.4 | 0.4 | 0.01 | 0.3 | 0.4 | 0.6 |
| Set 2 | 40 | 15 | 0.1 | 4 | 0.4 | 0.2 | 0.2 | 0.2 | 0.01 | 0.3 | 0.6 | 0.6 |
| Set 3 | 40 | 15 | 0.1 | 4 | 0.2 | 0.2 | 0.2 | 0.2 | 0.01 | 0.3 | 0.6 | 0.6 |
| Set 4 | 25 | 15 | 2 | 2 | 0.4 | 0.4 | 0.6 | 0.6 | 0.4 | 0.2 | 0.6 | 0.6 |

**Table S4:** Kinetic parameters for the 121 voxels of whole cell system in Figs. 3a,e

|  | Intra-region |  |  |  |  |  |  |  |  | Inter-region |  |  |
| --- | --- | --- | --- | --- | --- | --- | --- | --- | --- | --- | --- | --- |
| | $k$ | $K$ | $\lambda$ | $d_M$ | $d_P$ | $D_M^n$ | $D_P^n$ | $D_M^c$ | $D_P^c$ | $D_M^{nc}$ | $D_P^{nc}$ | $D_P^{cn}$ |
| Value | 40 | 15 | 4 | 0.4 | 0.1 | 0.1 | 0.2 | 0.2 | 0.3 | 0.2 | 0.2 | 0.2 |

